# Enhancing Neurocognitive Outcome Prediction in Congenital Heart Disease Patients: The Role of Brain Age Biomarkers and Beyond

**DOI:** 10.1101/2023.09.01.555976

**Authors:** Mohammad Arafat Hussain, Ellen Grant, Yangming Ou

## Abstract

This paper aimed to investigate the predictive power of combining demographic, socioeconomic, and genetic factors with a brain MRI-based quantified measure of accelerated brain aging (referred to as *deltaAGE*) for neurocognitive outcomes in adolescents and young adults with Congenital Heart Disease (CHD). Our hypothesis posited that including the brain age biomarker (*deltaAGE*) would enhance neurocognitive outcome predictions compared to models excluding it. We conducted comprehensive analyses, including leave-one-subject-out and leave-one-group-out cross-validation techniques. Our results demonstrated that the inclusion of *deltaAGE* consistently improved prediction performance when considering the Pearson correlation coefficient, a preferable metric for this study. Notably, the *deltaAGE* -augmented models consistently outperformed those without *deltaAGE* across all cross-validation setups, and these correlations were statistically significant (*p*-value < 0.05). Therefore, our hypothesis that incorporating the brain-age biomarker alongside demographic, socioeconomic, and genetic factors enhances neurocognitive outcome predictions in adolescents and young adults with CHD is supported by the findings.

## 1 Introduction

Congenital heart disease (CHD) is the most common birth defect, occurring in 1% of live births^1, 2^, with an 85% survival rate into adulthood^3–7^. As a result, in the USA alone, *∼*30,000 infants with CHD survive per year with a normal life expectancy. However, *∼*50% of survivors develop neurodevelopmental impairments that emerge in adolescence^8–13^ through young adulthood^8, 14–17^. In fact, in the past 5 years studies have shown increasing concern^17–19^ for accelerated brain aging with increased risk of dementia in adolescence and young adulthood (8-30 years)^13, 20^.

Predicting neurocognitive outcomes is an urgent unmet need. In the USA, more than 30,000 infants per year survive CHD but half of them develop neurocognitive impairments in adulthood^8–17^. Intervention before adulthood may improve outcomes^21^, by promoting parent-child relationships, individual psycho-education, outreach to community healthcare providers^22^, and home-administered computerized training programs^23, 24^. However, the bottleneck problem is: how to identify CHD patients at high risk for neurocognitive impairments. High-risk patients should be ideal candidates for interventions, while interventions on low-risk patients should be avoided. Current work inferring adulthood neurocognitive outcomes are very few and mostly used clinical variables or traditional brain MRI metrics, i.e., volumes^14, 25^. However, they explain only 1/3 of the neurocognitive outcomes^26–28^, insufficient to support interventional clinical trials.

Early prediction of later-life neurocognitive outcomes will create a precious time window for early intervention^29, 30^. It will identify high-risk patients for targeted intervention, avoiding unnecessary interventions for patients at low risk for future neurocognitive impairment^31^. Both the early and the targeted interventions are key unmet needs in clinical trials that aim to improve CHD patients’ long-term neurocognitive outcomes^21, 30^. Very few existing studies use brain magnetic resonance images (MRIs), demographics, socioeconomic status (SES), or genetic factors to predict later-life neurocognitive outcomes^32–38^. In this paper, we test our overall hypothesis that combining demographics, SES, or genetic factors, and adding a brain MRI-based quantified severity of accelerated brain aging, can better predict neurocognitive outcomes than without the brain age biomarker.

## 2 Methods

### 2.1 Data

We accessed the brain MRI and the associated demographic, SES, and genetic data of 89 patients from the Pediatric Cardiac Genomics Consortium (PCGC) database. The institutional review board of the Boston Children’s Hospital approved the access to data (Approval numbers IRB P00039087 and P00023574). In Table 1, we present the collection site, demographic, socioeconomic, genetic, and diagnosis details of this dataset. Further, in Table 2, we show the performed neurocognitive tests on the patients in this dataset. In this study, we use different prediction models to predict only those scores, which are available for all 89 subjects. We also used age, sex, diagnostic group, and data collection sites as independent variables and standardized all the scores before training and validating our prediction models.

**Table 1.**
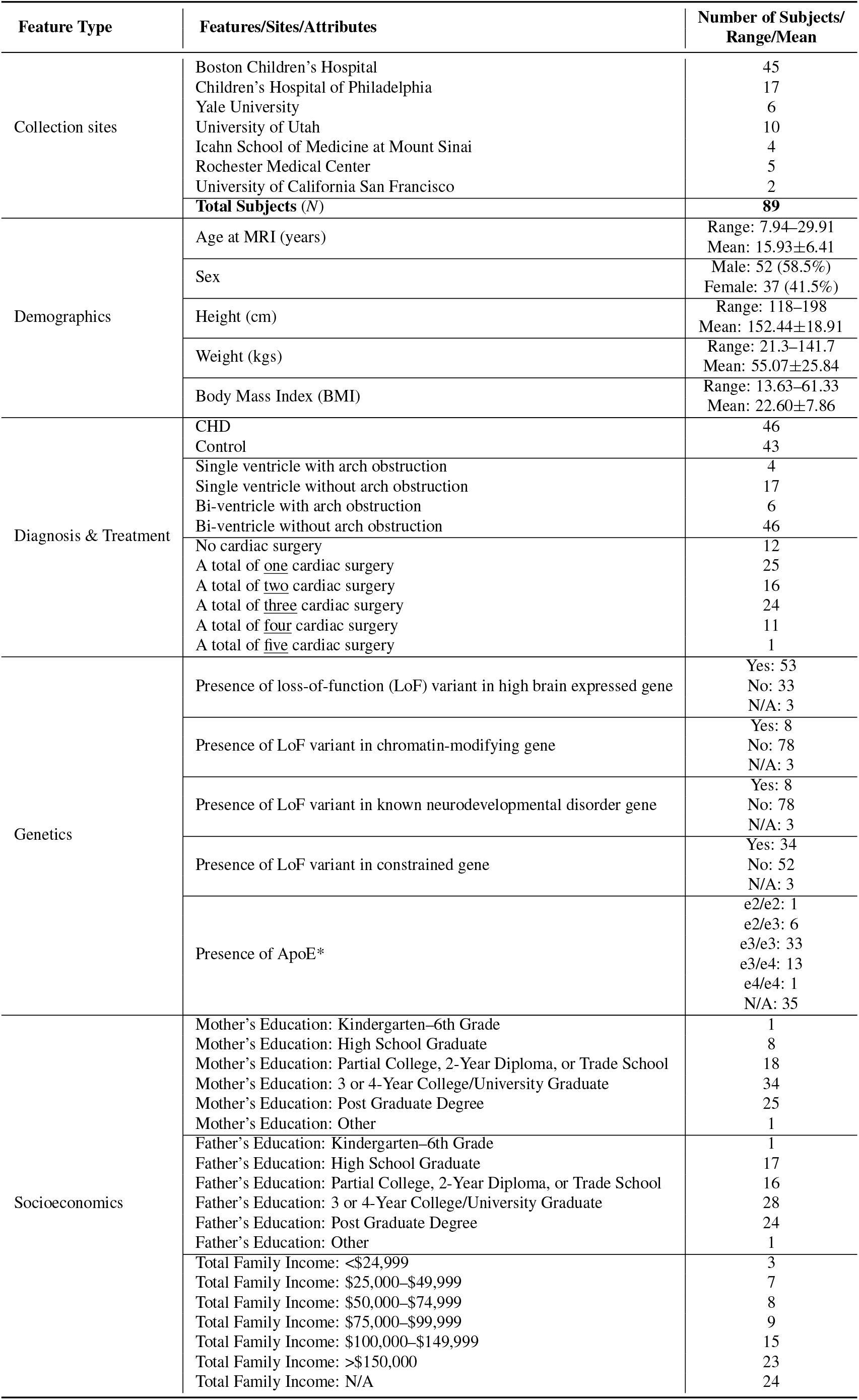
Demographic, socioeconomic, genetic, diagnosis, treatment, and collection site details of the 89 patients used in this study. (*Not used as a feature in the present study)

**Table 2.** A list of neurocognitive tests associated with the PCGC data, and the number of patients with scores available per test. (*Not used as a target score for prediction in the present study)

| Test Name | Number of Subjects |
| --- | --- |
| Word Reading | 89 |
| Sentence Comprehension | 89 |
| Spelling | 89 |
| Math Computation | 89 |
| Reading Composite | 89 |
| Block Design | 89 |
| Similarities | 89 |
| Digit Span | 89 |
| Matrix Reasoning | 89 |
| Vocabulary | 89 |
| Arithmetic* | 43 |
| Symbol Search | 89 |
| Visual Puzzle* | 43 |
| Information Extraction* | 43 |
| Coding | 89 |
| Verbal Comprehension Index | 89 |
| Perceptual Reasoning Index* | 43 |
| Working Memory Index* | 43 |
| Processing Speed Index | 89 |
| Full-Scale IQ | 89 |
| Fluid Reasoning Index* | 46 |
| Figure Weights* | 46 |

### 2.2 Neurocognitive Score Prediction

To test the hypothesis that combining brain MRI-based quantified severity of accelerated brain aging to demographics, SES, and genetic factors can better predict neurocognitive scores than without the brain-age biomarker, we train and validate each of our predictive models in leave-one-sample-out as well as leave-one-group-out cross-validation in two stages. In the first stage, we use all features including the brain-age biomarker, and in the second stage, we use all features except the brain-age biomarker.

#### 2.2.1 Estimation of Accelerated Brain Aging

In a recent study^39^, we trained a deep learning brain age estimator on T1-weighted brain MRIs of 16,705 healthy brain MRIs^40^ and produced the prediction of brain age for 96 adolescents and young adult survivors of CHD (accessed from the same PCGC dataset as in this study; 8-30 years of age). We computed the severity of brain aging by subtracting deep model-estimated and the actual chronological ages (*deltaAGE*; we refer to it as the MRI-based brain-age biomarker henceforth). Using a T-test with normal controls, we confirmed the existence and severity of accelerated brain aging (i.e., *deltaAGE* > 0 with *p*-value < 0.05).

#### 2.2.2 Prediction by Regression Forests

We used a regression forest that ensembles regression decisions from 5-layered (i.e., depth) 100 decision trees (see Fig. 1(a)). As described earlier, in the first stage of leave-one-sample-out and leave-one-group-out cross-validation, we used *deltaAGE* along with other features mentioned in Table 1 to predict neurocognitive scores mentioned in Table 2. Note that some of the features or target scores are not available for many patients and that is why, we did not include those features and target scores in the present study (see Tables 1 and 2). In the second stage of leave-one-sample-out and leave-one-group-out cross-validation, we train the regression forests again from scratch using the same features as used in stage one, except *deltaAGE*, to predict neurocognitive scores. Also note that in both stages, we train our regression forest from scratch for each of the neurocognitive scores separately.

**Figure 1.**
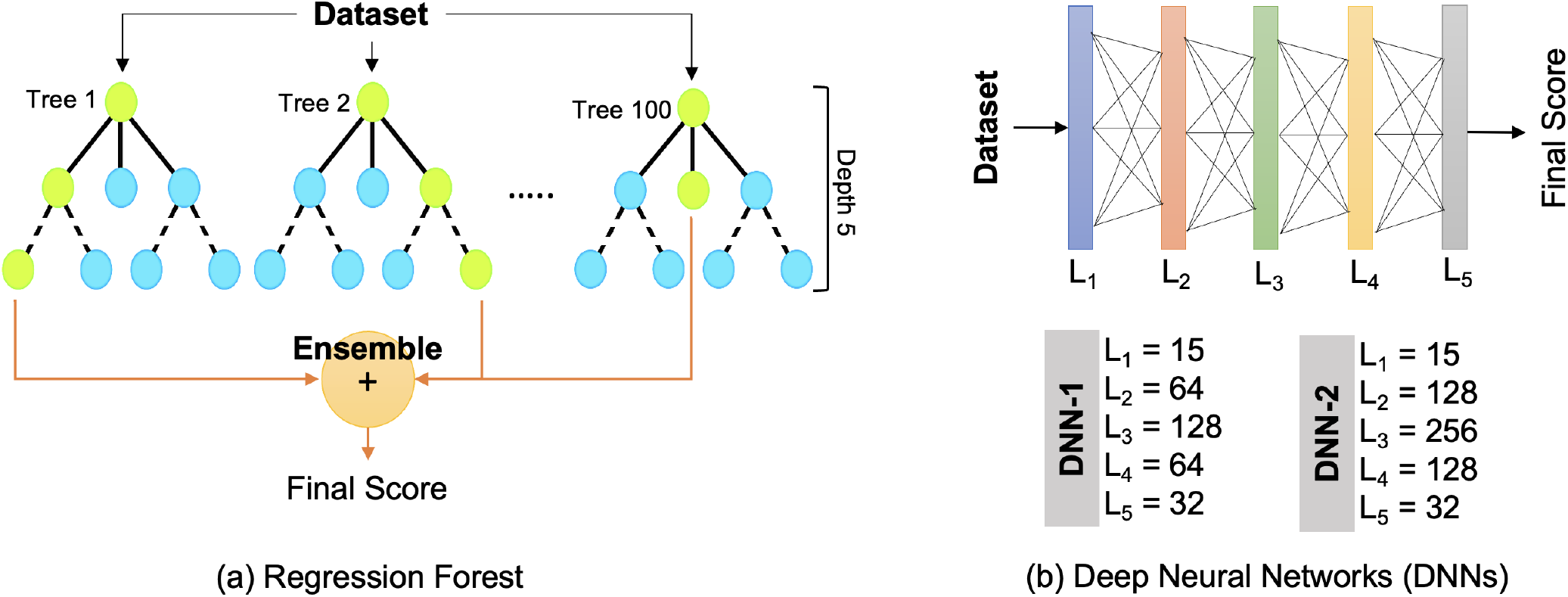
Prediction models in this study. (a) Regression forest that ensembles regression decisions from 100 regression trees, each of depth five, and (b) 5-layer deep neural networks (DNNs) of two settings. The number of nodes in each hidden fully connected layer is also shown for each DNN setting. DNN-2 is wider than DNN-1.

#### 2.2.3 Prediction by DNNs

We also used two 5-layered DNNs of different widths (i.e., different numbers of nodes in the hidden layers) in leave-one-sample-out and leave-one-group-out cross-validation setups. Like regression forests, in the first stage, we used *deltaAGE* along with other features mentioned in Table 1 to train DNNs to predict neurocognitive scores mentioned in Table 2. In the second stage, we train those DNNs again from scratch using the same features as used in stage one, except *deltaAGE*, to predict neurocognitive scores. We used the mean absolute error loss function to train these DNNs defined as:

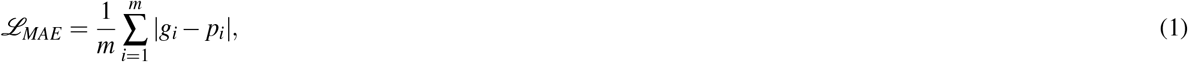

where *g* and *p* are the ground truth and the predicted test scores, respectively, and *m* denotes the total number of training data in a batch. We chose the Adam optimizer with a learning rate of 0.001 to train both DNNs. We also chose a batch size of 16. We implemented our models in PyTorch version 1.6.0 and Python version 3.8.10. The training was performed on the E2 cluster of Boston Children’s Hospital using an Intel E5-2650 v4 Broadwell 2.2 GHz processor, an Nvidia Titan RTX GPU with 24 GB of VRAM, and 8 GB of RAM.

### 2.3 Metrics for Prediction Accuracy Evaluation

To evaluate the neurocognition prediction accuracy, we used the Pearson correlation coefficient (*r*), mean absolute error (MAE), and mean absolute percentage error (MAPE) between the predicted and the ground-truth scores. The Pearson correlation between paired datasets (*G, P*) : *{*(*g*_1_, *p*_1_), (*g*_2_, *p*_2_), …, (*g*_*N*_, *p*_*N*_)*}* is mathematically defined as^41^:

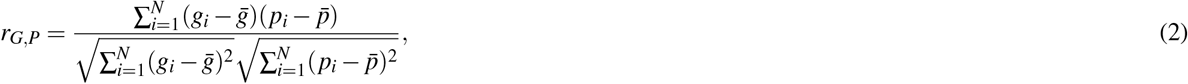

where 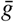 and 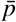 are the mean of all data points in the ground truth and predicted scores *G* and *P*, respectively, and *N* denotes the total number of test data. In addition, the MAE metric is defined as:

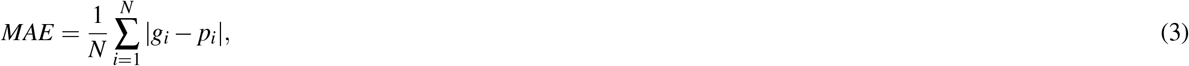

where *g* and *p* are the ground truth and the predicted values, respectively, and *N* denotes the total number of test data. Furthermore, the MAPE metric is defined as:

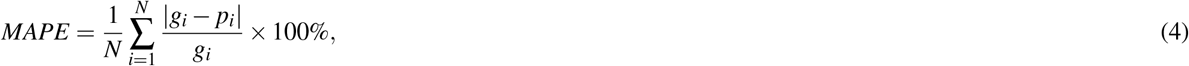

where *g* and *p* are the ground truth and the predicted values, respectively, and *N* denotes the total number of test data.

## 3 Results

In this section, we provide comparative neurocognitive score prediction performance by the regression forest, DNN-1, and DNN-2 in terms of Pearson correlation, MAE, MAPE, and the Wilcoxon signed-rank in leave-on-sample-out and leave-one-group-out cross-validation setup. Further, we used the data collection sites as well as the diagnosis, i.e., either CHD or control cohort, as groups. We ultimately compared this performance using two feature sets, once with *deltaAGE* and again without *deltaAGE*.

### 3.1 Leave-one-sample-out Performance

In Tables 3 and 4, we show the leave-one-subject-out cross-validated prediction performance by the regression forest, DNN-1, and DNN-2. We show the Pearson correlation coefficient (*r*) between the actual and predicted neurocognitive test scores in Table 3, where we present two sets of results by the regression forest, DNN-1, and DNN-2 for each neurocognitive test. For one set, we combined the brain-age bio-marker (i.e., *deltaAGE*) with other features (shown in columns with header ‘w/ *deltaAGE*’ in tables), while for another set we did not combine *deltaAGE* with other features (shown in columns with header ‘w/o *deltaAGE*’ in tables). We see in Table 3 that prediction performance by the regression forest is overall better than that by the DNN-1 and DNN-2, and the correlation (*r*) is statistically significant (for *p*-value=0.05) for many tests. On the other hand, the correlation between the actual scores and DNN-predicted scores is worse and not statistically significant (for *p*-value=0.05) for any test. Therefore, relying more on regression forest-based prediction, we further see that the prediction performance is better in the occasion when *deltaAGE* is combined with other features as confirmed by the higher mean of correlation coefficient (see first column under regression forest in Table 3).

**Table 3.**
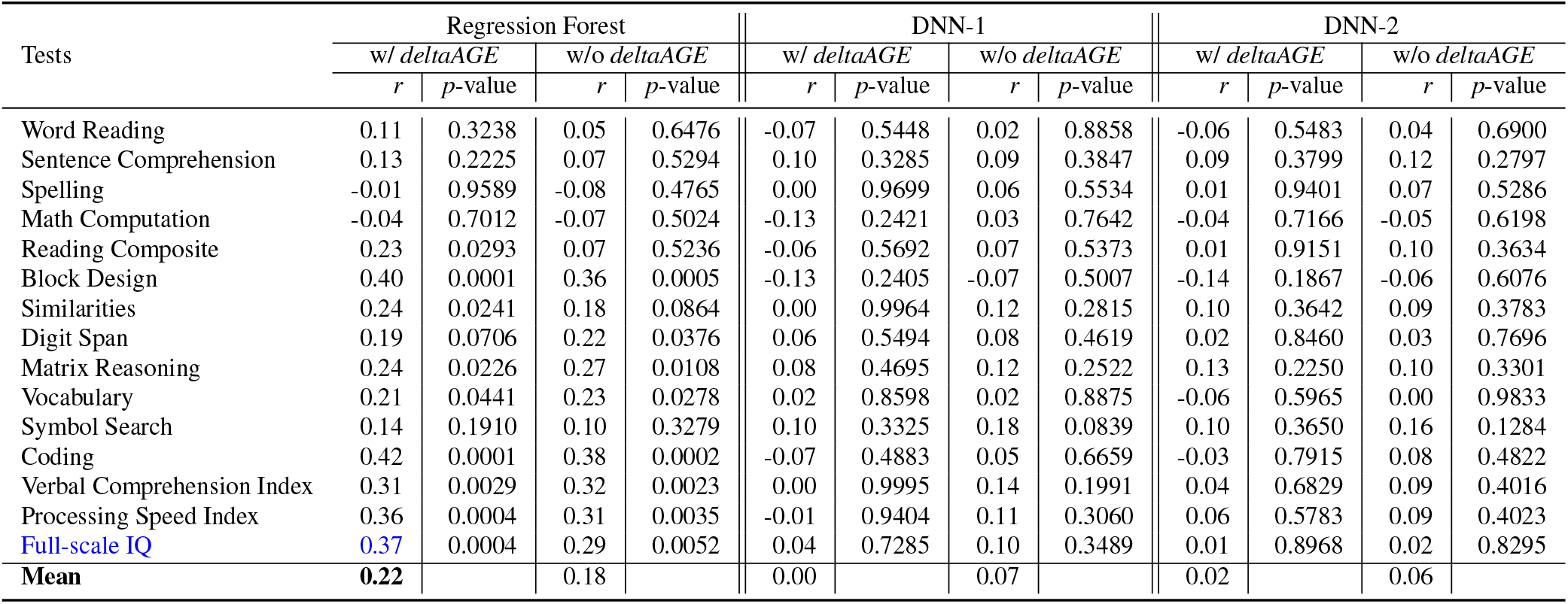
Pearson correlation performance between the ground truth and predicted neurocognitive scores by the regression forest, DNN-1, and DNN-2 in leave-one-sample-out cross-validation. The best correlation value for full-scale IQ is shown in blue font and the best mean correlation value is shown in **bold** font. Acronyms-w/: with, w/o: without, IQ: intelligent quotient.

| Tests | Regression Forest |  |  |  | DNN-1 |  |  |  | DNN-2 |  |  |  |
| --- | --- | --- | --- | --- | --- | --- | --- | --- | --- | --- | --- | --- |
|  | w/ *deltaAGE* |  | w/o *deltaAGE* |  | w/ *deltaAGE* |  | w/o *deltaAGE* |  | w/ *deltaAGE* |  | w/o *deltaAGE* |  |
|  | *r* | *p*-value | *r* | *p*-value | *r* | *p*-value | *r* | *p*-value | *r* | *p*-value | *r* | *p*-value |
| Word Reading | 0.11 | 0.3238 | 0.05 | 0.6476 | -0.07 | 0.5448 | 0.02 | 0.8858 | -0.06 | 0.5483 | 0.04 | 0.6900 |
| Sentence Comprehension | 0.13 | 0.2225 | 0.07 | 0.5294 | 0.10 | 0.3285 | 0.09 | 0.3847 | 0.09 | 0.3799 | 0.12 | 0.2797 |
| Spelling | -0.01 | 0.9589 | -0.08 | 0.4765 | 0.00 | 0.9699 | 0.06 | 0.5534 | 0.01 | 0.9401 | 0.07 | 0.5286 |
| Math Computation | -0.04 | 0.7012 | -0.07 | 0.5024 | -0.13 | 0.2421 | 0.03 | 0.7642 | -0.04 | 0.7166 | -0.05 | 0.6198 |
| Reading Composite | 0.23 | 0.0293 | 0.07 | 0.5236 | -0.06 | 0.5692 | 0.07 | 0.5373 | 0.01 | 0.9151 | 0.10 | 0.3634 |
| Block Design | 0.40 | 0.0001 | 0.36 | 0.0005 | -0.13 | 0.2405 | -0.07 | 0.5007 | -0.14 | 0.1867 | -0.06 | 0.6076 |
| Similarities | 0.24 | 0.0241 | 0.18 | 0.0864 | 0.00 | 0.9964 | 0.12 | 0.2815 | 0.10 | 0.3642 | 0.09 | 0.3783 |
| Digit Span | 0.19 | 0.0706 | 0.22 | 0.0376 | 0.06 | 0.5494 | 0.08 | 0.4619 | 0.02 | 0.8460 | 0.03 | 0.7696 |
| Matrix Reasoning | 0.24 | 0.0226 | 0.27 | 0.0108 | 0.08 | 0.4695 | 0.12 | 0.2522 | 0.13 | 0.2250 | 0.10 | 0.3301 |
| Vocabulary | 0.21 | 0.0441 | 0.23 | 0.0278 | 0.02 | 0.8598 | 0.02 | 0.8875 | -0.06 | 0.5965 | 0.00 | 0.9833 |
| Symbol Search | 0.14 | 0.1910 | 0.10 | 0.3279 | 0.10 | 0.3325 | 0.18 | 0.0839 | 0.10 | 0.3650 | 0.16 | 0.1284 |
| Coding | 0.42 | 0.0001 | 0.38 | 0.0002 | -0.07 | 0.4883 | 0.05 | 0.6659 | -0.03 | 0.7915 | 0.08 | 0.4822 |
| Verbal Comprehension Index | 0.31 | 0.0029 | 0.32 | 0.0023 | 0.00 | 0.9995 | 0.14 | 0.1991 | 0.04 | 0.6829 | 0.09 | 0.4016 |
| Processing Speed Index | 0.36 | 0.0004 | 0.31 | 0.0035 | -0.01 | 0.9404 | 0.11 | 0.3060 | 0.06 | 0.5783 | 0.09 | 0.4023 |
| Full-scale IQ | **0.37** | 0.0004 | 0.29 | 0.0052 | 0.04 | 0.7285 | 0.10 | 0.3489 | 0.01 | 0.8968 | 0.02 | 0.8295 |
| Mean | **0.22** |  | 0.18 |  | 0.00 |  | 0.07 |  | 0.02 |  | 0.06 |  |

We also show the MAE and MAPE performance between the actual and predicted neurocognitive test scores for ‘with *deltaAGE*’ and ‘without *deltaAGE*’ by the regression forest, DNN-1, and DNN-2 for each neurocognitive test in Table 4. Further, we estimated the difference between the actual and predicted scores for ‘with *deltaAGE*’ and ‘without *deltaAGE*’ cases followed by the Wilcoxon signed-rank test. The Wilcoxon signed-rank test produces a statistic value of 0 when two distributions perfectly match to each other. Otherwise, the statistic value gets larger as the two distribution gets further away from each other. We see in Table 4 that prediction performance in terms of the MAE and MAPE by the regression forest is overall better than that by the DNN-1 and DNN-2 as depicted by the least mean of MAE and MAPE by the regression forest (see first and second columns under regression forest in Table 4). Further, we see that the Wilcoxon signed-rank statistic value is large between the ‘with *deltaAGE*’ and ‘without *deltaAGE*’ cases for regression forest, which infer that prediction performance for ‘with *deltaAGE*’ is better than that for the ‘without *deltaAGE*’ case, although this statistic value of not statistically significant (for *p*-value=0.05).

**Table 4.** MAE and MAPE performance, and the Wilcoxon signed-rank test between the ground truth and predicted neurocognitive scores by the regression forest, DNN-1, and DNN-2 in leave-on-sample-out cross-validation setup. The least MAE and MAPE values are shown in **bold** font. Acronyms-w/: with, w/o: without, C: comprehension, I: index, IQ: intelligent quotient, W. Stat.: Wilcoxon signed-rank statistic.

| Tests | Regression Forest |  |  |  |  |  | DNN-1 |  |  |  |  |  | DNN-2 |  |  |  |  |  |
| --- | --- | --- | --- | --- | --- | --- | --- | --- | --- | --- | --- | --- | --- | --- | --- | --- | --- | --- |
|  | w/ *deltaAGE* |  | w/o *deltaAGE* |  | w/ vs. w/o *deltaAGE* |  | w/ *deltaAGE* |  | w/o *deltaAGE* |  | w/ vs. w/o *deltaAGE* |  | w/ *deltaAGE* |  | w/o *deltaAGE* |  | w/ vs. w/o *deltaAGE* |  |
|  | MAE | MAPE | MAE | MAPE | W. Stat. | p-value | MAE | MAPE | MAE | MAPE | W. Stat. | p-value | MAE | MAPE | MAE | MAPE | W. Stat. | p-value |
| Word Reading | 10.64 | 0.10 | 11.07 | 0.11 | 1650.00 | 0.817 | 14.98 | 0.14 | 13.88 | 0.13 | 1881.00 | 0.493 | 16.15 | 0.16 | 15.27 | 0.15 | 1810.0 | 0.431 |
| Sentence C | 13.76 | 0.26 | 13.98 | 0.26 | 1732.00 | 0.170 | 17.54 | 0.29 | 15.96 | 0.28 | 1719.00 | 0.006 | 16.56 | 0.29 | 16.71 | 0.29 | 1639.0 | 0.137 |
| Spelling | 11.99 | 0.23 | 12.68 | 0.23 | 1952.00 | 0.859 | 15.62 | 0.26 | 15.34 | 0.26 | 1590.00 | 0.135 | 16.17 | 0.27 | 15.70 | 0.26 | 1873.0 | 0.596 |
| Math Computation | 13.70 | 0.15 | 13.66 | 0.15 | 1969.00 | 0.966 | 17.94 | 0.19 | 16.29 | 0.17 | 1917.00 | 0.255 | 16.48 | 0.17 | 17.24 | 0.18 | 1968.0 | 0.888 |
| Reading Composite | 10.55 | 0.10 | 11.24 | 0.11 | 1874.00 | 0.898 | 15.77 | 0.15 | 15.04 | 0.15 | 1855.00 | 0.305 | 15.57 | 0.15 | 15.38 | 0.15 | 1832.0 | 0.485 |
| Block Design | 2.15 | 0.27 | 2.19 | 0.28 | 1899.00 | 0.678 | 2.50 | 0.33 | 2.50 | 0.33 | 1683.00 | 0.000 | 2.62 | 0.35 | 2.53 | 0.34 | 1659.0 | 0.160 |
| Similarities | 2.60 | 0.32 | 2.68 | 0.32 | 1989.00 | 0.329 | 2.86 | 0.34 | 2.78 | 0.33 | 1903.00 | 0.443 | 2.88 | 0.34 | 2.84 | 0.34 | 1837.0 | 0.498 |
| Digit Span | 2.26 | 0.25 | 2.22 | 0.24 | 1685.00 | 0.979 | 2.46 | 0.27 | 2.40 | 0.26 | 1921.00 | 0.175 | 2.64 | 0.29 | 2.41 | 0.26 | 1807.0 | 0.424 |
| Matrix Reasoning | 2.07 | 0.22 | 2.10 | 0.22 | 1833.00 | 0.901 | 2.55 | 0.26 | 2.34 | 0.25 | 1819.00 | 0.274 | 2.43 | 0.26 | 2.45 | 0.26 | 1914.0 | 0.717 |
| Vocabulary | 2.23 | 0.23 | 2.25 | 0.23 | 1902.00 | 0.321 | 2.62 | 0.28 | 2.61 | 0.27 | 1927.00 | 0.002 | 2.61 | 0.28 | 2.70 | 0.28 | 1898.0 | 0.669 |
| Symbol Search | 2.02 | 0.24 | 2.07 | 0.24 | 1923.00 | 0.292 | 2.14 | 0.25 | 2.15 | 0.25 | 1940.00 | 0.158 | 2.24 | 0.26 | 2.19 | 0.25 | 1917.0 | 0.727 |
| Coding | 1.87 | 0.24 | 1.89 | 0.25 | 1698.00 | 0.436 | 2.38 | 0.32 | 2.40 | 0.31 | 1768.00 | 0.294 | 2.33 | 0.30 | 2.28 | 0.30 | 1304.0 | 0.004 |
| Verbal CI | 12.66 | 0.12 | 12.74 | 0.12 | 1919.00 | 0.705 | 16.81 | 0.16 | 16.30 | 0.16 | 1746.00 | 0.596 | 16.51 | 0.16 | 16.76 | 0.16 | 1682.0 | 0.190 |
| Processing Speed I | 9.33 | 0.10 | 9.52 | 0.10 | 1808.00 | 0.038 | 14.68 | 0.15 | 13.15 | 0.14 | 1548.00 | 0.128 | 14.61 | 0.15 | 13.85 | 0.14 | 1786.0 | 0.376 |
| Full-scale IQ | 10.95 | 0.11 | 10.92 | 0.11 | 1739.00 | 0.619 | 14.44 | 0.15 | 15.35 | 0.15 | 1535.00 | 0.363 | 15.26 | 0.16 | 16.35 | 0.16 | 1924.0 | 0.748 |
| **Mean** | **7.25** | **0.20** | 7.41 | 0.21 | 1838.13 |  | 9.69 | 0.24 | 9.23 | 0.23 | 1783.47 |  | 9.67 | 0.24 | 9.64 | 0.24 | 1790.0 |  |

### 3.2 Leave-one-group-out Performance

We further tested the neurocognitive score prediction performance by the regression forest, DNN-1, and DNN-2 in leave-one-group-out cross-validation. We considered two different attributes for two different group-based cross-validation. First, we used data collection sites as groups, and second, diagnosis (i.e., CHD and control cohorts) as groups. In the following sections, we present those findings.

#### 3.2.1 Cohorts of Seven Data Collection Sites as Groups

In Tables 5 and 6, we show the leave-one-group-out cross-validated prediction performance by the regression forest, DNN-1, and DNN-2 for ‘with *deltaAGE*’ and ‘without *deltaAGE*’ cases. We show the Pearson correlation coefficient (*r*) between the actual and predicted neurocognitive test scores in Table 5, where see that prediction performance by the regression forest is overall better than that by the DNN-1 and DNN-2, and the correlation (*r*) statistically significant (for *p*-value=0.05) for several tests including ‘Full-scale IQ.’ On the other hand, the correlation between the actual scores and DNN-predicted scores is worse and not statistically significant (for *p*-value=0.05) for any test. Furthermore, we see that the prediction performance by the regression forest is better for the ‘with *deltaAGE*’ than the ‘without *deltaAGE*’ case (see first column under ‘Regression Forest’ in Table 5).

**Table 5.** Pearson correlation performance between the ground truth and predicted neurocognitive scores by the regression forest, DNN-1, and DNN-2 in leave-one-group-out cross-validation. In this table, we use data collection sites as groups. The best correlation value for full-scale IQ is shown in blue font and the best mean correlation value is shown in **bold** font. Acronyms-w/: with, w/o: without, IQ: intelligent quotient.

| Tests | Regression Forest |  |  |  | DNN-1 |  |  |  | DNN-2 |  |  |  |
| --- | --- | --- | --- | --- | --- | --- | --- | --- | --- | --- | --- | --- |
|  | w/ *deltaAGE* |  | w/o *deltaAGE* |  | w/ *deltaAGE* |  | w/o *deltaAGE* |  | w/ *deltaAGE* |  | w/o *deltaAGE* |  |
|  | *r* | *p*-value | *r* | *p*-value | *r* | *p*-value | *r* | *p*-value | *r* | *p*-value | *r* | *p*-value |
| Word Reading | 0.21 | 0.0443 | 0.20 | 0.0620 | -0.01 | 0.9588 | 0.05 | 0.6438 | -0.05 | 0.6201 | 0.03 | 0.7670 |
| Sentence Comprehension | 0.11 | 0.2938 | 0.08 | 0.4628 | 0.16 | 0.1251 | 0.17 | 0.1158 | 0.07 | 0.5308 | 0.18 | 0.0955 |
| Spelling | 0.09 | 0.3863 | 0.06 | 0.5571 | 0.07 | 0.5236 | 0.06 | 0.5687 | 0.02 | 0.8620 | 0.09 | 0.4011 |
| Math Computation | 0.06 | 0.5824 | 0.06 | 0.5932 | -0.05 | 0.6567 | 0.03 | 0.7746 | -0.04 | 0.6914 | 0.02 | 0.8644 |
| Reading Composite | 0.26 | 0.0131 | 0.13 | 0.2119 | 0.11 | 0.3167 | 0.10 | 0.3503 | 0.07 | 0.5152 | 0.07 | 0.5278 |
| Block Design | 0.17 | 0.1173 | 0.14 | 0.1940 | -0.12 | 0.2742 | -0.15 | 0.1749 | -0.13 | 0.2308 | -0.15 | 0.1648 |
| Similarities | 0.24 | 0.0217 | 0.17 | 0.1164 | 0.02 | 0.8399 | 0.08 | 0.4572 | 0.05 | 0.6707 | 0.12 | 0.2588 |
| Digit Span | 0.24 | 0.0223 | 0.25 | 0.0188 | 0.01 | 0.9176 | 0.11 | 0.3185 | -0.02 | 0.8239 | 0.04 | 0.6906 |
| Matrix Reasoning | 0.13 | 0.2333 | 0.11 | 0.2955 | 0.00 | 0.9665 | 0.07 | 0.4925 | 0.13 | 0.2297 | 0.07 | 0.5042 |
| Vocabulary | 0.08 | 0.4376 | 0.13 | 0.2126 | 0.11 | 0.3214 | 0.01 | 0.9051 | 0.03 | 0.8105 | 0.07 | 0.5087 |
| Symbol Search | 0.07 | 0.4965 | 0.05 | 0.6492 | 0.06 | 0.5733 | 0.13 | 0.2258 | 0.04 | 0.7127 | 0.11 | 0.2856 |
| Coding | 0.13 | 0.2165 | 0.26 | 0.0125 | -0.06 | 0.5650 | -0.02 | 0.8391 | -0.02 | 0.8580 | 0.05 | 0.6395 |
| Verbal Comprehension Index | 0.19 | 0.0745 | 0.21 | 0.0481 | 0.10 | 0.3532 | 0.07 | 0.5291 | 0.05 | 0.6147 | 0.11 | 0.3241 |
| Processing Speed Index | 0.15 | 0.1593 | 0.22 | 0.0394 | 0.08 | 0.4689 | 0.15 | 0.1657 | 0.07 | 0.5097 | 0.04 | 0.7197 |
| Full-scale IQ | **0.27** | 0.0096 | 0.25 | 0.0175 | 0.09 | 0.4235 | 0.08 | 0.4486 | 0.06 | 0.6005 | 0.09 | 0.3888 |
| **Mean** | **0.16** |  | 0.15 |  | 0.04 |  | 0.06 |  | 0.02 |  | 0.06 |  |

We further show the MAE and MAPE performance between the actual and predicted neurocognitive test scores for ‘with *deltaAGE*’ and ‘without *deltaAGE*’ by the regression forest, DNN-1, and DNN-2 for each neurocognitive test in Table 6. We also estimated the difference between the actual and predicted scores for ‘with *deltaAGE*’ and ‘without *deltaAGE*’ cases followed by the Wilcoxon signed-rank test. We see in Table 6 that prediction performance in terms of the MAE and MAPE by the regression forest is overall better than that by the DNN-1 and DNN-2 as depicted by the least mean of MAE and MAPE by the regression forest (see first and second columns under ‘Regression Forest’ in Table 6). Further, we see that the Wilcoxon signed-rank statistic value is large between the ‘with *deltaAGE*’ and ‘without *deltaAGE*’ cases for regression forest, which infer that prediction performance for ‘with *deltaAGE*’ is better than that for the ‘without *deltaAGE*’ case, although this statistic value of not statistically significant (for *p*-value=0.05).

**Table 6.**
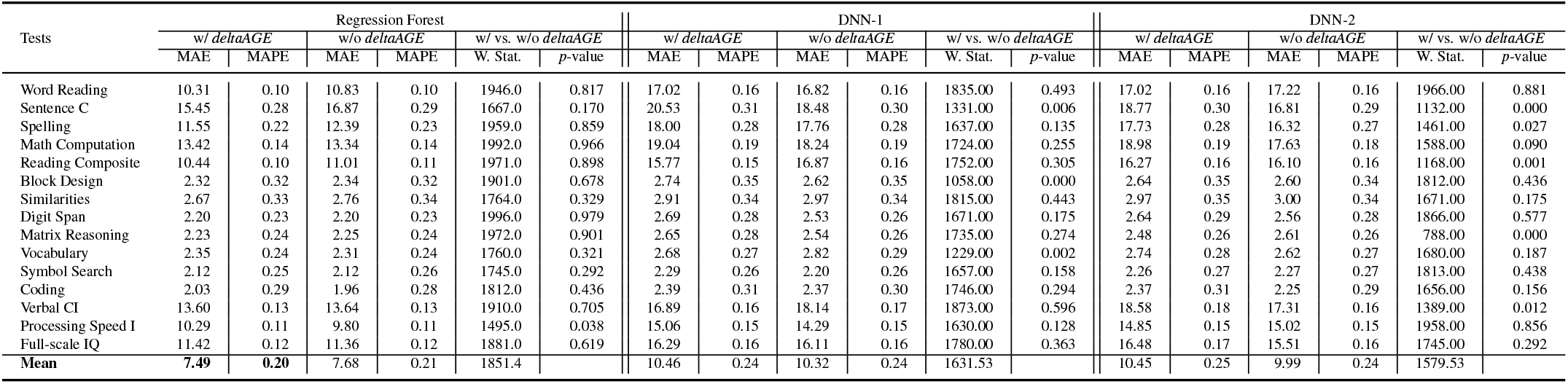
MAE and MAPE performance, and the Wilcoxon signed-rank test between the ground truth and predicted neurocognitive scores by the regression forest, DNN-1, and DNN-2 in leave-one-group-out cross-validation. In this table, we use data collection sites as groups. The least MAE and MAPE values are shown in **bold** font. Acronyms-w/: with, w/o: without, C: comprehension, I: index, IQ: intelligent quotient, W. Stat.: Wilcoxon signed-rank statistic.

#### 3.2.2 CHD and Control Cohorts as Groups

In Tables 7, 8, and 9, we show the leave-one-group-out cross-validated prediction performance in terms of Pearson correlation coefficient (*r*) between the actual and predicted neurocognitive test scores by the regression forest, DNN-1, and DNN-2, respectively, for ‘with *deltaAGE*’ and ‘without *deltaAGE*’ cases. In these tables, we used the CHD and control cohorts as groups. We see in these tables that prediction performance by all the approaches (i.e., regression forest, DNN-1, and DNN-2) are found to be better for ‘without *deltaAGE*’ case and when control cohort is used for training and CHD cohort for validation, as depicted by the best mean *r* in respective tables of regression forests, DNN-1, and DNN-2 (see the second last columns under ‘LOGO (Training: Control, Test: CHD)’ in Tables 7, 8, and 9). In addition, the prediction performance in terms of *r* is found to be the best for the DNN-2 approach (see Table 9), although we see correlation performance to be statistically significant (for *p*-value=0.05) for fewer tests than we found for the regression forest in leave-one-sample-out cross-validations in Table 3, and leave-one-group-out (where, data collection sites as group) cross-validations in Table 5.

**Table 7.** Pearson correlation performance between the ground truth and predicted neurocognitive scores by the regression forest in leave-one-group-out (LOGO) cross-validation. In this table, we used the CHD cohort for training and the control cohort for validation, and vice versa. The best correlation value for full-scale IQ is shown in blue font and the best mean correlation value is shown in **bold** font. Acronyms-w/: with, w/o: without, IQ: intelligent quotient.

| Tests | LOGO (Training: CHD, Test: Control) |  |  |  | LOGO (Training: Control, Test: CHD) |  |  |  |
| --- | --- | --- | --- | --- | --- | --- | --- | --- |
|  | w/ *deltaAGE* |  | w/o *deltaAGE* |  | w/ *deltaAGE* |  | w/o *deltaAGE* |  |
|  | *r* | *p*-value | *r* | *p*-value | *r* | *p*-value | *r* | *p*-value |
| Word Reading | 0.08 | 0.6227 | 0.08 | 0.5932 | -0.16 | 0.2998 | 0.10 | 0.5028 |
| Sentence Comprehension | -0.12 | 0.4583 | -0.19 | 0.2103 | 0.06 | 0.6808 | -0.06 | 0.7071 |
| Spelling | -0.10 | 0.5126 | -0.17 | 0.2767 | -0.07 | 0.6394 | 0.04 | 0.8081 |
| Math Computation | 0.10 | 0.5141 | 0.09 | 0.5571 | 0.08 | 0.6102 | 0.23 | 0.1301 |
| Reading Composite | -0.06 | 0.6936 | 0.02 | 0.9137 | -0.06 | 0.6927 | 0.20 | 0.1752 |
| Block Design | 0.16 | 0.2959 | 0.14 | 0.3855 | 0.12 | 0.4149 | 0.17 | 0.2563 |
| Similarities | 0.10 | 0.5094 | 0.13 | 0.4121 | 0.11 | 0.4683 | 0.09 | 0.5423 |
| Digit Span | 0.10 | 0.5290 | 0.08 | 0.5973 | 0.17 | 0.2545 | 0.15 | 0.3171 |
| Matrix Reasoning | 0.19 | 0.2176 | 0.13 | 0.3957 | 0.11 | 0.4622 | 0.11 | 0.4869 |
| Vocabulary | -0.15 | 0.3426 | -0.05 | 0.7593 | 0.07 | 0.6451 | 0.12 | 0.4315 |
| Symbol Search | 0.16 | 0.3121 | 0.09 | 0.5607 | 0.21 | 0.1533 | 0.17 | 0.2705 |
| Coding | 0.42 | 0.0051 | 0.38 | 0.0130 | 0.29 | 0.0515 | 0.38 | 0.0091 |
| Verbal Comprehension Index | 0.10 | 0.5424 | 0.08 | 0.5987 | 0.15 | 0.3232 | 0.16 | 0.2894 |
| Processing Speed Index | 0.32 | 0.0353 | 0.24 | 0.1200 | 0.36 | 0.0127 | 0.33 | 0.0240 |
| Full-scale IQ | 0.19 | 0.2129 | 0.04 | 0.7896 | 0.13 | 0.3974 | 0.17 | 0.2564 |
| Mean | 0.10 |  | 0.07 |  | 0.10 |  | **0.16** |  |

**Table 8.** Pearson correlation performance between the ground truth and predicted neurocognitive scores by the DNN-1 in leave-one-group-out (LOGO) cross-validation. In this table, we used the CHD cohort for training and the control cohort for validation, and vice versa. The best correlation value for full-scale IQ is shown in blue font and the best mean correlation value is shown in **bold** font. Acronyms-w/: with, w/o: without, IQ: intelligent quotient.

| Tests | LOGO (Training: CHD, Test: Control) |  |  |  | LOGO (Training: Control, Test: CHD) |  |  |  |
| --- | --- | --- | --- | --- | --- | --- | --- | --- |
|  | w/ *deltaAGE* |  | w/o *deltaAGE* |  | w/ *deltaAGE* |  | w/o *deltaAGE* |  |
|  | *r* | *p*-value | *r* | *p*-value | *r* | *p*-value | *r* | *p*-value |
| Word Reading | -0.29 | 0.0573 | -0.24 | 0.1154 | 0.13 | 0.3866 | 0.18 | 0.2340 |
| Sentence Comprehension | -0.04 | 0.8117 | -0.21 | 0.1854 | 0.33 | 0.0242 | 0.38 | 0.0083 |
| Spelling | -0.15 | 0.3381 | -0.03 | 0.8720 | 0.14 | 0.3635 | 0.24 | 0.1114 |
| Math Computation | -0.32 | 0.0380 | -0.24 | 0.1193 | 0.21 | 0.1619 | 0.19 | 0.2143 |
| Reading Composite | -0.26 | 0.0964 | -0.18 | 0.2563 | 0.31 | 0.0380 | 0.32 | 0.0324 |
| Block Design | -0.39 | 0.0089 | -0.28 | 0.0668 | 0.01 | 0.9335 | 0.09 | 0.5740 |
| Similarities | -0.28 | 0.0686 | -0.03 | 0.8676 | 0.25 | 0.0889 | 0.22 | 0.1506 |
| Digit Span | -0.24 | 0.1208 | -0.02 | 0.9088 | 0.06 | 0.7125 | 0.22 | 0.1428 |
| Matrix Reasoning | -0.11 | 0.4781 | -0.16 | 0.3144 | 0.19 | 0.2068 | 0.20 | 0.1834 |
| Vocabulary | -0.10 | 0.5065 | -0.20 | 0.2030 | 0.33 | 0.0272 | 0.29 | 0.0515 |
| Symbol Search | 0.03 | 0.8413 | 0.12 | 0.4400 | 0.12 | 0.4349 | 0.17 | 0.2722 |
| Coding | 0.00 | 0.9839 | 0.05 | 0.7366 | 0.17 | 0.2675 | 0.30 | 0.0441 |
| Verbal Comprehension Index | -0.10 | 0.5048 | -0.01 | 0.9636 | 0.34 | 0.0215 | 0.27 | 0.0732 |
| Processing Speed Index | 0.10 | 0.5030 | 0.09 | 0.5566 | 0.06 | 0.7001 | 0.21 | 0.1530 |
| Full-scale IQ | -0.24 | 0.1143 | -0.16 | 0.3120 | 0.31 | 0.0362 | 0.20 | 0.1750 |
| Mean | -0.16 |  | -0.10 |  | 0.20 |  | **0.23** |  |

**Table 9.** Pearson correlation performance between the ground truth and predicted neurocognitive scores by the DNN-2 in leave-one-group-out (LOGO) cross-validation. In this table, we used the CHD cohort for training and the control cohort for validation, and vice versa. The best correlation value for full-scale IQ is shown in blue font and the best mean correlation value is shown in **bold** font. Acronyms-w/: with, w/o: without, IQ: intelligent quotient.

| Tests | LOGO (Training: CHD, Test: Control) |  |  |  | LOGO (Training: Control, Test: CHD) |  |  |  |
| --- | --- | --- | --- | --- | --- | --- | --- | --- |
| | w/ ΔAGE | | w/o ΔAGE | | w/ ΔAGE | | w/o ΔAGE | |
|  | *r* | *p*-value | *r* | *p*-value | *r* | *p*-value | *r* | *p*-value |
| Word Reading | -0.37 | 0.0157 | -0.18 | 0.2552 | 0.17 | 0.2665 | 0.18 | 0.2412 |
| Sentence Comprehension | -0.22 | 0.1607 | -0.10 | 0.5189 | 0.36 | 0.0151 | 0.33 | 0.0267 |
| Spelling | -0.12 | 0.4310 | 0.04 | 0.7930 | 0.18 | 0.2306 | 0.16 | 0.2829 |
| Math Computation | -0.32 | 0.0370 | -0.23 | 0.1343 | 0.14 | 0.3609 | 0.20 | 0.1909 |
| Reading Composite | -0.35 | 0.0202 | -0.23 | 0.1405 | 0.35 | 0.0179 | 0.36 | 0.0138 |
| Block Design | -0.41 | 0.0058 | -0.38 | 0.0109 | -0.05 | 0.7590 | 0.09 | 0.5403 |
| Similarities | -0.21 | 0.1872 | -0.12 | 0.4451 | 0.30 | 0.0426 | 0.23 | 0.1208 |
| Digit Span | -0.17 | 0.2739 | -0.12 | 0.4308 | 0.17 | 0.2489 | 0.25 | 0.0942 |
| Matrix Reasoning | -0.17 | 0.2786 | -0.03 | 0.8632 | 0.18 | 0.2443 | 0.27 | 0.0676 |
| Vocabulary | -0.22 | 0.1489 | -0.15 | 0.3384 | 0.23 | 0.1206 | 0.36 | 0.0154 |
| Symbol Search | 0.03 | 0.8508 | 0.16 | 0.3207 | 0.07 | 0.6650 | 0.22 | 0.1506 |
| Coding | -0.09 | 0.5670 | -0.06 | 0.7223 | 0.06 | 0.7106 | 0.16 | 0.2865 |
| Verbal Comprehension Index | -0.14 | 0.3653 | -0.21 | 0.1752 | 0.28 | 0.0563 | 0.37 | 0.0122 |
| Processing Speed Index | 0.00 | 0.9937 | 0.03 | 0.8710 | 0.23 | 0.1320 | 0.22 | 0.1491 |
| Full-scale IQ | -0.23 | 0.1428 | -0.11 | 0.4656 | **0.29** | 0.0475 | 0.27 | 0.0670 |
| **Mean** | -0.20 |  | -0.11 |  | 0.20 |  | **0.24** |  |

We also show the MAE and MAPE performance between the actual and predicted neurocognitive test scores for ‘with *deltaAGE*’ and ‘without *deltaAGE*’ by the regression forest, DNN-1, and DNN-2 for each neurocognitive test in Tables 10, 11, and 12, respectively. We further estimated the difference between the actual and predicted scores for ‘with *deltaAGE*’ and ‘without *deltaAGE*’ cases followed by the Wilcoxon signed-rank test and showed the Wilcoxon statistic and associated *p*-value in Tables 10, 11, and 12. We see in Table 6 that prediction performance in terms of the MAE and MAPE by the regression forest is overall better for ‘without *deltaAGE*’ case and when the control cohort is used for training and CHD cohort for validation, as depicted by the least mean MAE and mean MAPE. On the other hand, for DNN-1 and DNN-2, prediction performance in terms of the MAE and MAPE is overall better (i.e., least MAE and MAPE) for ‘without *deltaAGE*’ case but when CHD cohort is used for training and control cohort for validation (Tables 11 and 12). Thus, irrespective of the training and validation group, all approaches, i.e., regression forest, DNN-1, and DNN-2, performed better in prediction for the ‘without *deltaAGE*’ case. This is the opposite finding of what we found in the leave-one-subject-out and leave-one-group-out (the group being the data collection site) cross-validation setup as seen in Tables 4 and 6. Further, we see that the Wilcoxon signed-rank statistic value is smaller (< 500) between the ‘with *deltaAGE*’ and ‘without *deltaAGE*’ cases for regression forest, DNN-1, and DNN-2, compared to those (> 1500) for leave-one-subject-out and leave-one-group-out (group being data collection site) cross-validation setup as seen in Tables 4 and 6. It infers that prediction distributions for ‘with *deltaAGE*’ and ‘without *deltaAGE*’ are closer to each other when the leave-one-group-out setup uses CHD and control cohorts as groups.

**Table 10.** MAE and MAPE performance, and the Wilcoxon signed-rank test between the ground truth and predicted neurocognitive scores by the regression forest in leave-one-group-out (LOGO) cross-validation. In this table, we used the CHD cohort for training and the control cohort for validation, and vice versa. The least MAE and MAPE values are shown in **bold** font. Acronyms-w/: with, w/o: without, IQ: intelligent quotient.

| Tests | LOGO (Training: CHD, Test: Control) |  |  |  |  |  | LOGO (Training: Control, Test: CHD) |  |  |  |  |  |
| --- | --- | --- | --- | --- | --- | --- | --- | --- | --- | --- | --- | --- |
| | w/ ΔAGE | | w/o ΔAGE | | w/ vs. w/o ΔAGE | | w/ ΔAGE | | w/o ΔAGE | | w/ vs. w/o ΔAGE | |
|  | MAE | MAPE | MAE | MAPE | W. Stat. | *p*-value | MAE | MAPE | MAE | MAPE | W. Stat. | *p*-value |
| Word Reading | 11.1711 | 0.1127 | 11.8116 | 0.1186 | 419.0 | 0.5220 | 11.8432 | 0.1071 | 11.1981 | 0.1012 | 354.0 | 0.0414 |
| Sentence Comprehension | 13.6002 | 0.2266 | 13.8749 | 0.2297 | 463.0 | 0.9096 | 13.6533 | 0.2972 | 13.5617 | 0.3061 | 501.0 | 0.6729 |
| Spelling | 14.0151 | 0.3776 | 14.5038 | 0.3811 | 407.0 | 0.4328 | 14.7320 | 0.1344 | 15.2182 | 0.1402 | 410.0 | 0.1569 |
| Math Computation | 13.9749 | 0.1402 | 13.9858 | 0.1421 | 257.0 | 0.0083 | 13.3565 | 0.1468 | 12.2133 | 0.1325 | 268.0 | 0.0024 |
| Reading Composite | 12.0509 | 0.1255 | 11.8614 | 0.1235 | 462.0 | 0.9001 | 11.2439 | 0.1022 | 10.3844 | 0.0935 | 337.0 | 0.0256 |
| Block Design | 2.1823 | 0.2384 | 2.2843 | 0.2507 | 331.0 | 0.0876 | 2.5841 | 0.3922 | 2.5206 | 0.3851 | 510.0 | 0.7454 |
| Similarities | 3.0422 | 0.4195 | 2.9494 | 0.4146 | 355.0 | 0.1574 | 2.7351 | 0.2650 | 2.8378 | 0.2721 | 421.0 | 0.1955 |
| Digit Span | 2.6753 | 0.3329 | 2.7397 | 0.3410 | 333.0 | 0.0923 | 2.5100 | 0.2262 | 2.5614 | 0.2295 | 458.0 | 0.3738 |
| Matrix Reasoning | 1.9443 | 0.2040 | 2.0412 | 0.2152 | 462.0 | 0.9001 | 2.5741 | 0.2655 | 2.5778 | 0.2655 | 450.0 | 0.3287 |
| Vocabulary | 2.7102 | 0.3222 | 2.6630 | 0.3161 | 411.0 | 0.4616 | 2.2484 | 0.1857 | 2.2284 | 0.1840 | 367.0 | 0.0583 |
| Symbol Search | 1.7971 | 0.2173 | 1.9004 | 0.2295 | 415.0 | 0.4913 | 2.2447 | 0.2619 | 2.2244 | 0.2536 | 310.0 | 0.0110 |
| Coding | 1.7037 | 0.2440 | 1.7452 | 0.2517 | 448.0 | 0.7696 | 2.1192 | 0.2674 | 2.1094 | 0.2558 | 233.0 | 0.0005 |
| Verbal Comprehension Index | 15.0922 | 0.1497 | 15.2359 | 0.1526 | 259.0 | 0.0090 | 13.6165 | 0.1274 | 13.2727 | 0.1239 | 293.0 | 0.0062 |
| Processing Speed Index | 8.6544 | 0.0922 | 9.1939 | 0.0983 | 459.0 | 0.8718 | 10.0753 | 0.1070 | 10.4144 | 0.1091 | 247.0 | 0.0010 |
| Full-scale IQ | 11.2125 | 0.1168 | 12.2658 | 0.1280 | 380.0 | 0.2670 | 12.6172 | 0.1262 | 12.2527 | 0.1210 | 244.0 | 0.0009 |
| **Mean** | 7.7217 | 0.2213 | 7.9370 | 0.2261 | 390.7 |  | 7.8769 | 0.2008 | **7.7050** | **0.1982** | 360.2 |  |

**Table 11.** MAE and MAPE performance, and the Wilcoxon signed-rank test between the ground truth and predicted neurocognitive scores by the DNN-1 in leave-one-group-out (LOGO) cross-validation. In this table, we used the CHD cohort for training and the control cohort for validation, and vice versa. The least MAE and MAPE values are shown in **bold** font. Acronyms-w/: with, w/o: without, IQ: intelligent quotient.

| Tests | LOGO (Training: CHD, Test: Control) |  |  |  |  |  | LOGO (Training: Control, Test: CHD) |  |  |  |  |  |
| --- | --- | --- | --- | --- | --- | --- | --- | --- | --- | --- | --- | --- |
| | w/ ΔAGE | | w/o ΔAGE | | w/ vs. w/o ΔAGE | | w/ ΔAGE | | w/o ΔAGE | | w/ vs. w/o ΔAGE | |
|  | MAE | MAPE | MAE | MAPE | W. Stat. | *p*-value | MAE | MAPE | MAE | MAPE | W. Stat. | *p*-value |
| Word Reading | 13.9479 | 0.1299 | 14.7447 | 0.1347 | 395.0 | 0.3530 | 19.8863 | 0.1907 | 18.7606 | 0.1797 | 529.0 | 0.9053 |
| Sentence Comprehension | 19.0063 | 0.1795 | 15.6457 | 0.1470 | 264.0 | 0.0108 | 21.9458 | 0.4388 | 21.1289 | 0.4355 | 418.0 | 0.1843 |
| Spelling | 17.0929 | 0.1586 | 15.7440 | 0.1441 | 412.0 | 0.4689 | 18.8546 | 0.4009 | 19.6379 | 0.4144 | 415.0 | 0.1737 |
| Math Computation | 15.0337 | 0.1464 | 15.7065 | 0.1560 | 322.0 | 0.0689 | 22.7792 | 0.2396 | 20.6095 | 0.2153 | 518.0 | 0.8118 |
| Reading Composite | 14.5614 | 0.1369 | 14.4271 | 0.1355 | 385.0 | 0.2939 | 16.8978 | 0.1619 | 19.1627 | 0.1885 | 342.0 | 0.0296 |
| Block Design | 2.3164 | 0.3265 | 2.2749 | 0.3428 | 191.0 | 0.0004 | 3.1320 | 0.3744 | 2.9371 | 0.3629 | 359.0 | 0.0473 |
| Similarities | 2.4924 | 0.2961 | 2.6091 | 0.3108 | 280.0 | 0.0190 | 3.2987 | 0.3808 | 3.3002 | 0.3692 | 462.0 | 0.3976 |
| Digit Span | 2.4252 | 0.2376 | 2.5185 | 0.2484 | 460.0 | 0.8813 | 2.9390 | 0.3227 | 2.5361 | 0.2699 | 364.0 | 0.0539 |
| Matrix Reasoning | 2.5280 | 0.2933 | 2.4866 | 0.2791 | 280.0 | 0.0190 | 2.7586 | 0.2583 | 2.5969 | 0.2406 | 474.0 | 0.4745 |
| Vocabulary | 2.3692 | 0.2398 | 2.2670 | 0.2370 | 337.0 | 0.1022 | 2.9678 | 0.2932 | 3.3282 | 0.3368 | 282.0 | 0.0041 |
| Symbol Search | 2.2018 | 0.2822 | 2.0124 | 0.2633 | 348.0 | 0.1338 | 2.3634 | 0.2436 | 2.3672 | 0.2496 | 514.0 | 0.7784 |
| Coding | 2.3310 | 0.3850 | 2.0173 | 0.3417 | 467.0 | 0.9476 | 2.4497 | 0.2447 | 2.7041 | 0.2659 | 360.0 | 0.0486 |
| Verbal Comprehension Index | 13.7220 | 0.1329 | 14.1592 | 0.1370 | 426.0 | 0.5780 | 19.8477 | 0.1864 | 21.8574 | 0.2051 | 522.0 | 0.8456 |
| Processing Speed Index | 14.8407 | 0.1588 | 11.8377 | 0.1289 | 333.0 | 0.0923 | 15.2727 | 0.1478 | 16.5728 | 0.1609 | 530.0 | 0.9138 |
| Full-scale IQ | 13.7428 | 0.1417 | 12.5480 | 0.1307 | 382.0 | 0.2776 | 18.6767 | 0.1821 | 19.4363 | 0.1871 | 511.0 | 0.7536 |
| **Mean** | 9.2407 | 0.2163 | **8.7332** | **0.2091** | 352.1 |  | 11.6046 | 0.2710 | 11.7957 | 0.2720 | 440 |  |

**Table 12.** MAE and MAPE performance, and the Wilcoxon signed-rank test between the ground truth and predicted neurocognitive scores by the DNN-2 in leave-one-group-out (LOGO) cross-validation. In this table, we used the CHD cohort for training and the control cohort for validation, and vice versa. The least MAE and MAPE values are shown in **bold** font. Acronyms-w/: with, w/o: without, IQ: intelligent quotient.

| Tests | LOGO (Training: CHD, Test: Control) |  |  |  |  |  | LOGO (Training: Control, Test: CHD) |  |  |  |  |  |
| --- | --- | --- | --- | --- | --- | --- | --- | --- | --- | --- | --- | --- |
| | w/ $\Delta AGE$ | | w/o $\Delta AGE$ | | w/ vs. w/o $\Delta AGE$ | | w/ $\Delta AGE$ | | w/o $\Delta AGE$ | | w/ vs. w/o $\Delta AGE$ | |
|  | MAE | MAPE | MAE | MAPE | W. Stat. | p-value | MAE | MAPE | MAE | MAPE | W. Stat. | p-value |
| Word Reading | 13.2943 | 0.1231 | 15.1261 | 0.1409 | 356.0 | 0.1610 | 20.5105 | 0.1965 | 19.1704 | 0.1840 | 424.0 | 0.2071 |
| Sentence Comprehension | 14.7697 | 0.1376 | 13.2474 | 0.1248 | 208.0 | 0.0010 | 22.5004 | 0.4599 | 20.1314 | 0.4411 | 365.0 | 0.0553 |
| Spelling | 16.2302 | 0.1512 | 13.3411 | 0.1229 | 266.0 | 0.0116 | 19.1321 | 0.4002 | 19.1003 | 0.4036 | 491.0 | 0.5957 |
| Math Computation | 14.8633 | 0.1455 | 13.9531 | 0.1388 | 301.0 | 0.0375 | 22.8232 | 0.2406 | 21.0758 | 0.2228 | 506.0 | 0.7129 |
| Reading Composite | 13.3242 | 0.1248 | 12.6260 | 0.1216 | 306.0 | 0.0436 | 19.0143 | 0.1840 | 19.3446 | 0.1889 | 286.0 | 0.0048 |
| Block Design | 2.2457 | 0.3266 | 2.2006 | 0.3233 | 467.0 | 0.9476 | 3.0129 | 0.3664 | 2.9769 | 0.3611 | 450.0 | 0.3287 |
| Similarities | 2.6116 | 0.3217 | 2.5551 | 0.3027 | 323.0 | 0.0708 | 3.3059 | 0.3760 | 3.4086 | 0.3775 | 479.0 | 0.5087 |
| Digit Span | 2.3792 | 0.2412 | 2.2749 | 0.2365 | 459.0 | 0.8718 | 2.8777 | 0.3269 | 2.8348 | 0.3148 | 448.0 | 0.3180 |
| Matrix Reasoning | 2.2831 | 0.2748 | 2.4360 | 0.2722 | 53.0 | 0.0001 | 2.6618 | 0.2469 | 2.7815 | 0.2549 | 389.0 | 0.0993 |
| Vocabulary | 2.2523 | 0.2464 | 1.9998 | 0.2089 | 216.0 | 0.0015 | 3.1952 | 0.3168 | 3.2069 | 0.3203 | 474.0 | 0.4745 |
| Symbol Search | 2.1770 | 0.2891 | 1.9019 | 0.2501 | 471.0 | 0.9857 | 2.3401 | 0.2507 | 2.6100 | 0.2811 | 454.0 | 0.3508 |
| Coding | 2.1010 | 0.3673 | 2.0252 | 0.3511 | 468.0 | 0.9571 | 2.6123 | 0.2645 | 2.4581 | 0.2421 | 353.0 | 0.0402 |
| Verbal Comprehension Index | 15.2878 | 0.1494 | 12.1968 | 0.1191 | 338.0 | 0.1048 | 21.6491 | 0.2015 | 22.0946 | 0.2043 | 365.0 | 0.0553 |
| Processing Speed Index | 12.6538 | 0.1390 | 12.6416 | 0.1365 | 418.0 | 0.5143 | 16.9028 | 0.1662 | 17.2474 | 0.1707 | 441.0 | 0.2823 |
| Full-scale IQ | 14.0394 | 0.1445 | 12.4544 | 0.1299 | 433.0 | 0.6367 | 18.7664 | 0.1842 | 18.3644 | 0.1814 | 462.0 | 0.3976 |
| **Mean** | 8.7008 | 0.2121 | **8.0653** | **0.1986** | 338.8 |  | 12.0869 | 0.2787 | 11.7870 | 0.2765 | 425.8 |  |

### 3.3 Full-scale IQ Prediction Performance

Full-scale IQ is typically used to assess human general intelligence, the fundamental ability that combines all subdomains of neurocognitive abilities^42, 43^. These subdomain abilities can be assessed via different neurocognitive tests, some of which are available with the PCGC dataset we used in this study (see Table 2). However, the full-scale IQ provides an overall quantification of a person’s general intelligence. Furthermore, full-scale IQ is not calculated by typical averaging of all subdomain scores, but rather by employing factor load analysis, resulting in heterogenous subdomain contributions to full-scale IQ^42^. Therefore, we also check the effect of brain-age bio-marker in the prediction of the full-scale IQ in this study. In Fig. 2, we show plots of the best predicted full-scale IQ scores in each of the three cross-validation setups. We observe in all three cross-validation setups in Figs. 2(a-c) that the prediction models performed better with statistical significance (*p*-value=0.05) when the feature set included *deltaAGE*, although overall performance in terms of mean correlation over all the test sometimes differed as seen Tables 7, 8, and 9.

**Figure 2.**
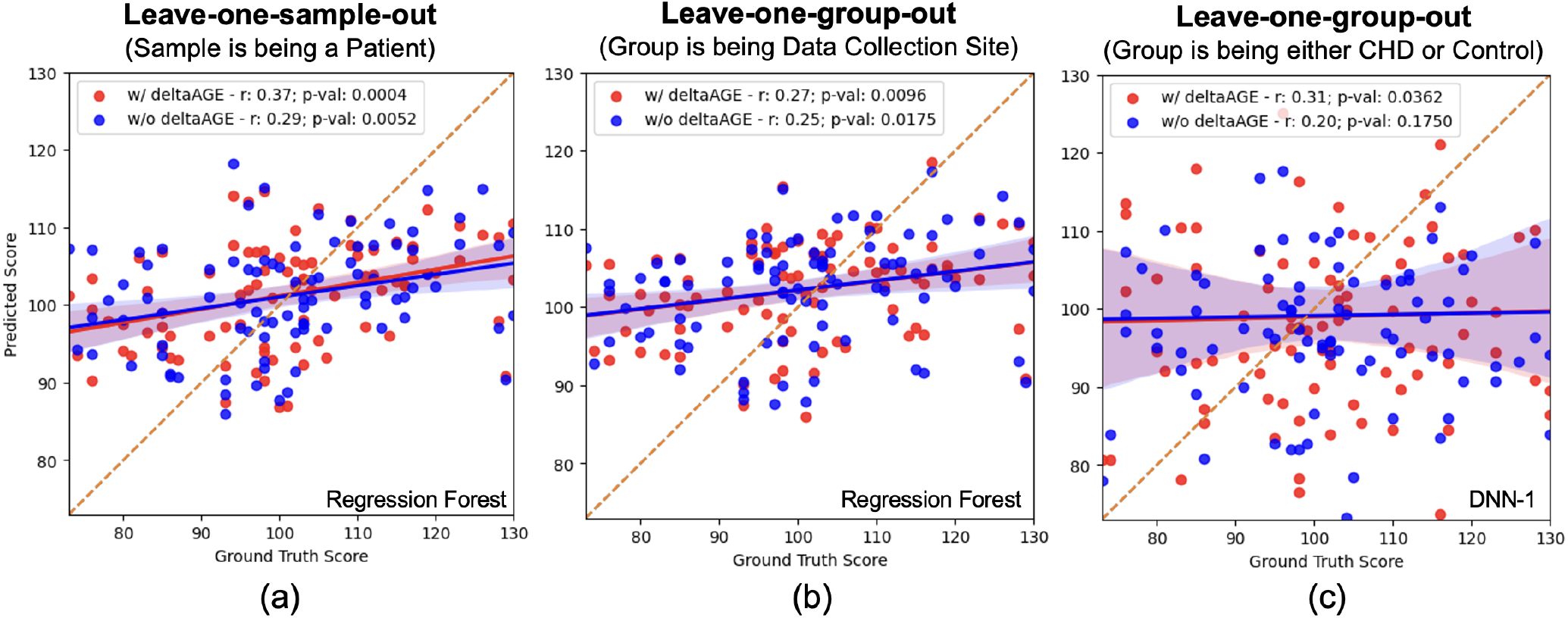
Best ‘Full-scale IQ’ score prediction performance in terms of Pearson correlation in three cross-validation setups. Best prediction (a) by the regression forest in the leave-one-sample-out cross-validation, (b) by the regression forest in leave-one-group-out cross-validation, where a group is the data collection site, and (c) by the DNN-1 in leave-one-group-out cross-validation, where a group being either CHD or control cohort.

## 4 Discussion

In this study, we conducted experiments to quantify the effect of brain-age bio-marker in predicting neurocognitive scores in adolescents and young adults with CHD. To perform this test, we employed regression forest and two DNNs on demographic, socioeconomic, and genetic factors. Our findings demonstrate the potential of leveraging MRI-based brain-age bio-marker for predicting neurocognition in patients with CHD. However, this investigation also presented us with several unanswered questions. For example, the size of the training data remains a concern when using deep neural network frameworks, as discussed by Richter et al.^44^. We had only 89 data samples in this study, which may not be sufficient to draw definitive conclusions. Despite these lingering questions, our study reveals that brain-age bio-marker can aid in predicting neurocognitive scores. Several key points arising from our findings warrant further discussion:

### 4.1 Pearson correlation *vs*. MAE

The Pearson correlation coefficient (*r*) seems more reliable than the MAE and MAPE metrics in neurocognition prediction accuracy estimation. The distribution of neurocognitive scores in our dataset follows a Gaussian-like distribution (see Fig. 3). As a result, a central tendency of the predicted scores towards the mean results in a low MAE and MAPE, although predictions were often inaccurate. Therefore, we considered *r* as a better indicator of the accuracy of the predicted IQ scores. In addition, the associated *p* value indicates the statistical significance of the estimated correlation.

**Figure 3.**
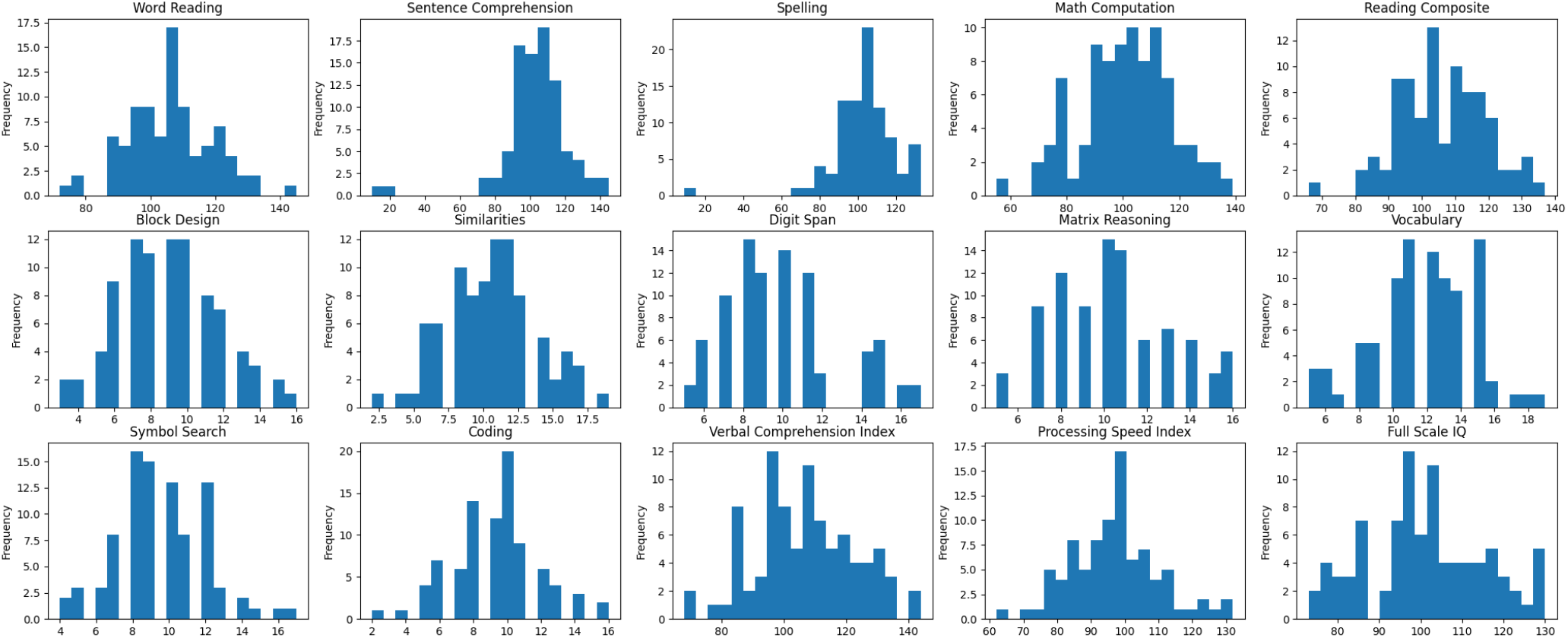
Ditribution of the ground truth scores of different neurocognitive tests.

### 4.2 Comparison to Structural MRI-based State-of-the-arts on Smaller Data

Several studies^45–47^ predicted the full-scale IQ score, which showed a correlation of 30-70% (*p <* 0.01) between the ground truth and estimated absolute full-scale IQ scores. These studies used a dataset of size less than 250 healthy subjects with an age distribution of 6–27 years. Our dataset consists of 89 patients with an age distribution of 7–30 years, and we achieved the best correlation between the actual and predicted full-scale IQ of 37% (see Table 3 and Fig. 2(a)) but without using structural MRI directly, rather employing demographic, socioeconomic, and genetic factors.

### 4.3 Test of Hypothesis

In this paper, we tested the hypothesis that combining demographics, socioeconomic, or genetic factors, and adding a brain MRI-based quantified severity of accelerated brain aging, can better predict neurocognitive outcomes than without the brain age biomarker. In our results, we observed that the mean correlation coefficient is better for the ‘with *deltaAGE*’ case than the ‘without *deltaAGE*’ in the leave-one-subject-out and the leave-one-group-out (where the group is the data collection site) as seen in Tables 3 and 5. However, in the leave-one-group-out (where the group is the CHD/control cohort) setup, the mean correlation coefficient is better for the ‘without *deltaAGE*’ case than the ‘with *deltaAGE*’ (see Tables 7, 8, and 9). Since,

1. The Pearson correlation is preferable to the MAE and MAPE metrics in this study (discussed earlier),
2. The size of the training data remains a concern when using DNN frameworks (discussed earlier),
3. Leave-one-group-out cross-validation with considering CHD/control cohorts as groups forced us to keep out about 50% of the data for validation, resulting in the smallest data size for training among our three cross-validation setup, and
4. Full-scale IQ is not calculated by typical averaging of all subdomain scores, but rather by employing factor load analysis, resulting in heterogenous subdomain contributions to full-scale IQ (discussed earlier),

We can consider Pearson correlation performance on full-scale IQ prediction by the leave-one-subject-out and the leave-one-group-out (where the group is the data collection site) more trustworthy than the overall mean correlation. Based on this consideration, we see in Fig. 2 that the Pearson correlation (*r*) for full-scale IQ prediction is better for ‘with *deltaAGE*’ than the ‘without *deltaAGE*’ in all three cross-validation setups. In addition, all the correlations between the ground truth and predicted full-scale IQ corresponding to ‘with *deltaAGE*’ cases are statistically significant (*p*-value < 0.05). Thus, our hypothesis that adding brain-age bio-marker to demographic, socioeconomic, and genetic factors in predicting neurocognition in adolescents and young adults with CHD stands true.

### 4.4 Limitations

While our paper presents valuable findings, it is important to acknowledge several limitations. Firstly, our study employed a relatively small sample size of only 89 patients. Increasing the sample size would enhance the statistical power and broaden the generalizability of our results. Secondly, by amalgamating data from both control and CHD groups, we may have introduced confounding variables, thereby restricting our ability to make specific conclusions about each group. Future investigations should contemplate analyzing these groups separately to gain a more precise understanding of the distinct contributions of brain structure to intelligence within each population. Furthermore, our analysis exclusively relied on a single dataset, potentially limiting the applicability of our findings to other populations or imaging protocols. To ensure the robustness of our results, it would be beneficial to validate them using multiple independent datasets. In addition, our study exclusively employed regression forest and DNN architectures and did not explore the potential advantages of utilizing alternative models such as support vector regressors or Vision Transformers (ViTs). Assessing various learning approaches could yield valuable insights and potentially enhance predictive performance. Lastly, our study concentrated solely on demographic, socioeconomic, genetic, and MRI-based brain-age biomarkers, omitting actual MRI features that could contribute to a more comprehensive understanding of the relationship between brain structure and intelligence. Future research should consider integrating these additional data sources to offer a more holistic perspective on intelligence prediction.

## Conclusion

In conclusion, this study provided valuable insights into the prediction of neurocognitive outcomes in CHD patients. Our results highlighted the utility of including a brain MRI-based bio-marker, *deltaAGE*, in predictive models, showing consistent improvements in prediction performance. However, it is essential to acknowledge several limitations, such as the relatively small sample size and the amalgamation of data from both control and CHD groups, potentially introducing confounding variables. Future research should address these limitations by increasing sample sizes, analyzing groups separately, and validating findings with multiple independent datasets. Moreover, exploring alternative machine learning models could offer further improvements in predictive accuracy. Additionally, integrating actual MRI features into the analysis could provide a more comprehensive understanding of the relationship between brain structure and intelligence. Overall, this study contributes to our understanding of neurocognitive prediction in CHD patients and paves the way for further research in this field.

## Acknowledgements

This work is supported by the American Heart Association grant (No. 919799).

